# The delivery of nano-formulated drugs to solid tumours is selectively increased by co-application of the vascular disrupting agent CA4P

**DOI:** 10.1101/2025.08.10.669501

**Authors:** Annabel Kitowski, Constanze Heise, Stefanie Sperling, Jasmin Hotz, Danica Bajic, Marina Rubey, Kathrin Klar, Georg Höfner, Julien Vollaire, Veronique Josserand, Jean-Luc Coll, Oliver Thorn-Seshold, Petar Marinković, Julia Thorn-Seshold

## Abstract

Improving the efficacy of existing cytotoxic chemotherapeutics requires increasing drug delivery to tumours while minimising systemic toxicity. Formulating these drugs as nanoparticles can reduce their exposure to healthy tissues, but broadly applicable strategies to enhance tumoral accumulation are lacking. Here, we show that co-administering small molecule vascular disrupting agents together with nanoparticle formulations (e.g. diagnostic reporters, or clinical drugs irinotecan and doxorubicin) increases their tumoral uptake by up to threefold, without raising systemic exposure. In a syngeneic mouse model of triple-negative breast cancer, this enhancement diminished when co-treatments were repeated, limiting its therapeutic benefit. However, since most solid tumour types are susceptible to vascular disrupting agents, this approach may be a broadly applicable strategy to improve the selectivity of drug delivery: with particular relevance for single dose use in diagnostic or research settings.

**Graphical Abstract:** 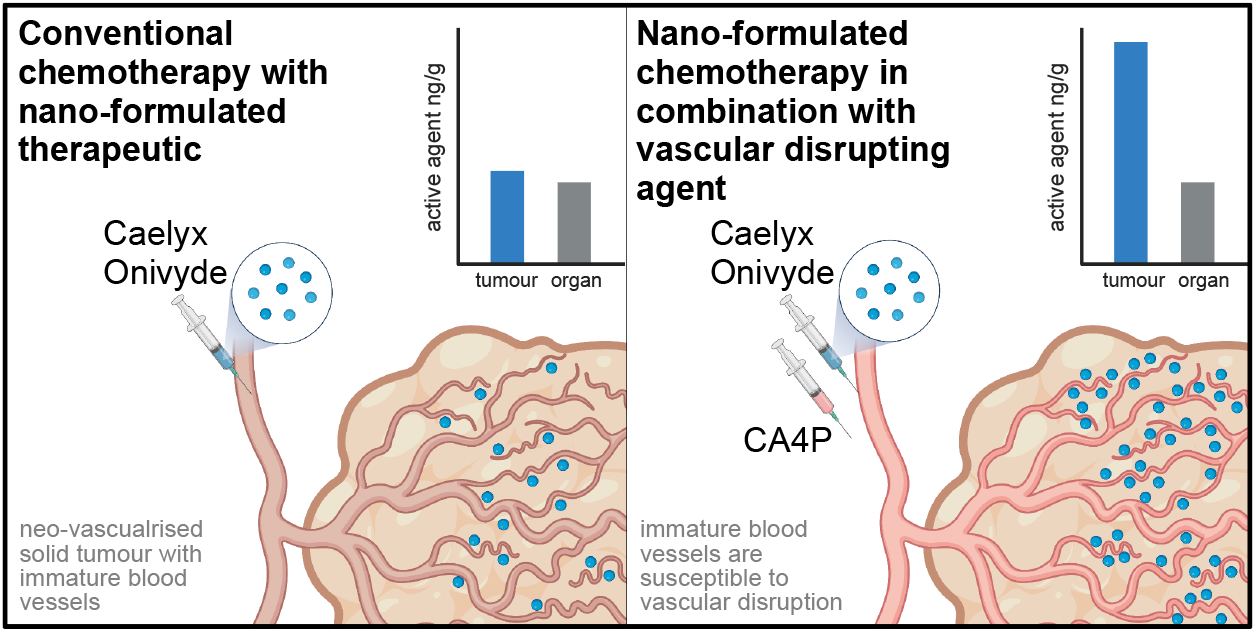

## Introduction

A challenge for all cytostatic chemotherapies is to achieve sufficient drug accumulation in tumours to ensure efficacy, while limiting systemic exposure to minimise toxicity. Liposomal nanoparticle formulations can limit systemic toxicity by enabling slow, controlled drug release, thereby improving tolerability compared to bolus administration of the free drugs (Liu et al., 2022). Several liposomal formulations are clinically approved, including for doxorubicin (Caelyx^®^, Janssen) and irinotecan (Onivyde^®^; Ipsen/Servier), and further nano-formulations are in advanced development (Miguel et al., 2022; Thapa & Kim, 2023).

However, broadly effective strategies to selectively enhance tumour delivery of nanoparticles are not known. The enhanced permeability and retention (EPR) effect, long considered the cornerstone of passive tumour targeting, has failed to translate consistently in clinical settings (Rosenblum et al., 2018; Sharifi et al., 2022). While personalised nanoparticles targeting tumour-specific antigens show promise (Chehelgerdi et al., 2023; Sun et al., 2023), their clinical utility is constrained by the cost and limited accessibility of detailed tumour profiling and production, which are rarely feasible outside research environments.

Here, we exploit a widely conserved hallmark of solid tumours: the low structural resilience of tumour-associated vasculature. Tumoral vessels grow abnormally, and do not develop the integrity of physiological vasculature. Small molecule vascular disrupting agents (VDAs) are drugs that selectively destabilise tumour vessels, by a range of mechanisms. The major class of VDAs are microtubule depolymerising drugs. When applied systemically, they rapidly destabilise microtubules in all endothelial cells. Healthy vasculature is mechanically stabilised by pericytes and ECM, so can maintain its integrity despite this insult; but tumoral vessels are not stabilised in this way, so they transiently collapse: an effect that is consistent across a large range of solid tumour types (Siemann, 2011; Tozer, Kanthou, et al., 2005; Tozer et al., 2008).

Microtubule-depolymerising VDAs such as combretastatin A4-phosphate (CA4P) and BNC105P were developed since the mid-1990s with the aim to use vascular shutdown to kill tumours by necrosis (Grisham et al., 2018; Prise et al., 2002). Despite effective core necrosis, Phase III clinical trials of VDA monotherapies failed (Seidi et al., 2017; Siemann & Horsman, 2009) since a viable tumour rim sustained by normal vasculature re-initiated tumour recurrence and malignant progression (Hinnen & Eskens, 2007; Liang et al., 2016).

In this work, we test whether VDAs can be used instead as “targeting agents” contributing their proven *tumour-selectivity*, but without requiring them to *also* be treatment effectors. We hypothesised that subtherapeutic doses of VDAs, that tumour-selectively slow blood flow and increase vessel permeability, could transiently enhance the extravasation of circulating nanoparticles by permitting longer and more effective interaction with a leakier vessel wall: thus tumour-selectively improving the delivery of nanoparticles or nano-formulated drugs. This could improve treatment outcomes with drugs that are already effective as monotherapies, but which suffer from side-effects due to effects in healthy tissues. As well as effects dependent on VDA dosage, we anticipated that the size of the nanoparticles, as well as the timing of the VDA co-administration, would affect the size and nature of changes to the tumoral delivery profile.

In mouse studies, we now show that VDA co-administration triples tumour accumulation of nano-encapsulated irinotecan, doxorubicin, and fluorescent reporter particles, without elevating systemic or off-target organ exposure. This enhancement diminishes with repeated VDA applications, limiting the therapeutic gain in our murine tumour models. However, these findings delineate a potentially generalisable strategy to augment tumour-selective nanoparticle delivery: with implications for both diagnostics and therapy, as well as basic research into the development of the complex microenvironment of solid tumours.

## Results

### VDA co-administration selectively enhances nanoparticle accumulation in solid tumours

To test whether vascular disruption can enhance nanoparticle delivery to tumours without affecting healthy tissue, we first conducted imaging-based biodistribution studies using NIR-fluorescent reporter nanoparticles (SAIVI, ThermoFisher). These PEGylated liposomal formulations of cyanine-type small molecule fluorophores allow both longitudinal *in vivo* imaging (qualitative), and endpoint quantification via fluorophore extraction from tissue. A subcutaneous, syngeneic murine triple-negative breast cancer model (4T1) was chosen. Expecting that VDA effects would most strongly influence the uptake of nanoparticles that are usually slow to penetrate to tumours, we screened a range of nanoparticle sizes (40-320 nm; **Figure S1**) while maintaining the same featureless PEGylated surface chemistry (no intrinsic targeting of nanoparticles to tumour). Animals were administered the reporter particles either alone, or together with a separate injection of the VDA combretastatin-A4 phosphate (CA4P). CA4P is typically used preclinically at 100 mg/kg dosage to totally cut off tumour blood flow (Dark et al., 1997); we instead applied far lower doses (10-30 mg/kg), aiming only to transiently permeabilise tumoral vasculature and reducing tumour blood flow rate (**Figure 1A**).

**Figure 1.**
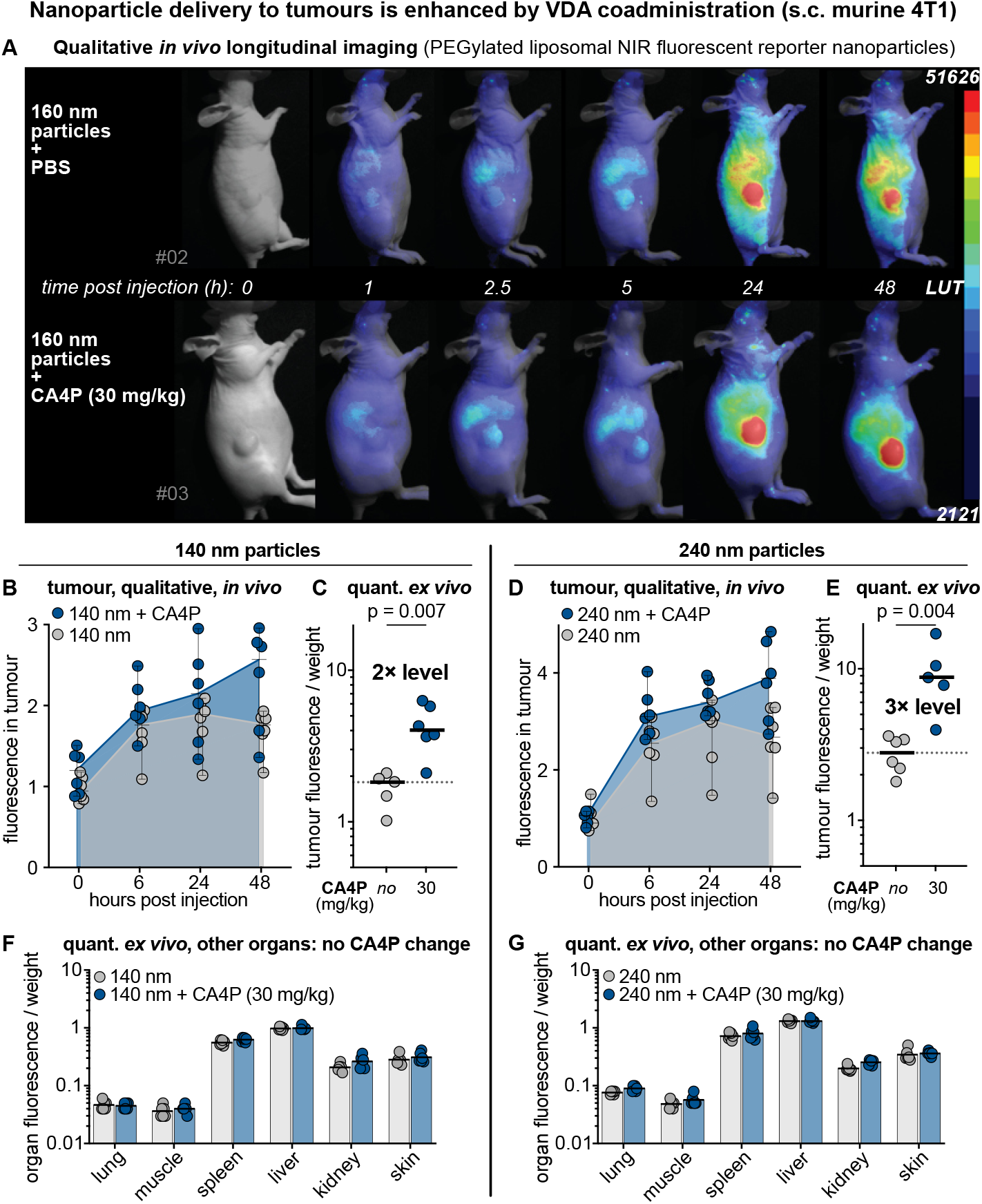
Transient vascular disruption enhances tumour-specific accumulation of fluorescent nanoparticles. (A) Representative whole-animal fluorescence images following intravenous injection of NIR-labelled reporter nanoparticles (SAIVI, ThermoFisher), administered with (bottom) or without (top) CA4P (related data: see **Figure S1**). (B, D) Longitudinal in vivo imaging of nanoparticle accumulation in subcutaneous 4T1 tumours for 140 nm (B) and 240 nm (D) particles. (C, E) Endpoint quantification of tumour fluorescence by fluorophore extraction from excised tumours, shown as fluorophore normalised to tumour tissue weight. Mann-Whitney U test. (G) Endpoint quantification of fluorescence in healthy organs (liver, kidney, lung, spleen, skin, muscle) shows no difference between groups, confirming the enhancements in panels C and E are tumour selective. Data shown as mean with 95% CI, n ≥ 5 mice per group.

Qualitative monitoring of tumour fluorescence over time by *in vivo* imaging indicated faster increase and higher endpoint levels of tumoral signal in CA4P-treated mice, for all nanoparticle sizes (**Figure 1B, D**; **Figure S1**). Endpoint analysis of extracted fluorophore per unit tissue weight quantified that the higher tumoral accumulation with CA4P is size-dependent and is maximised for nanoparticle sizes 140-240 nm (140 nm: >2-fold; 240 nm: 3-fold; **Figure 1C, E**). Importantly, for all nanoparticle sizes, the fluorophore levels in healthy organs (liver, kidney, lung, spleen, skin and muscle) remained unchanged (**Figure 1F, G**; **Figure S1**). Taken together, these data are consistent with the subtherapeutic VDA dose causing tumour-selective enhancement of nanoparticle delivery, without altering its systemic biodistribution.

### Co-administering CA4P improves delivery of nano-formulated chemotherapeutics to tumours without increasing exposure in healthy organs

We next tested whether the delivery of clinically used nano-formulations of chemotherapeutic drugs to tumours can be similarly selectively enhanced by VDA co-treatment, in an orthotopic 4T1 mouse model. Pegylated liposomal doxorubicin (Caelyx^®^, Baxter) and irinotecan (Onivyde^®^; Ipsen/Servier) are two frequently clinically used nano-formulations of long-approved, effective, cytostatic drugs that disrupt cell replication and cause cell death (irinotecan is metabolised to SN-38 which inhibits the enzyme topoisomerase I, while doxorubicin directly inhibits topoisomerase II). We examined their pharmacokinetics, with or without CA4P, in tumours and in healthy tissues, over 96 h after injection, by *ex vivo* drug extraction, followed by LCMS/MS quantification. These included metabolite analyses (since although doxorubicin is an active drug species, irinotecan is a pharmacologically inactive prodrug that must be converted *in situ* to the active agent SN-38 for efficacy), and isotope shift calibrants to control for extraction efficiencies (*Supporting Information* MS/MS).

#### Caelyx^®^ (doxorubicin)

Coadministration of CA4P (30 mg/kg) together with Caelyx^®^ (1 mg/kg) more than doubled cumulative doxorubicin exposure in tumours compared to Caelyx alone (area under the curve, AUC; **Figure 2A, B**). This effect was mainly driven by superior drug retention after 24 h. As a proxy measure for the risk of systemic side-effects, we also quantified doxorubicin in healthy tissues. We found no CA4P-dependent differences in heart and skin (two known toxicity sites in Caelyx therapy; **Figure 2C, D**), nor in plasma and lung (**Figure S2A, B**). With the VDA, tumoral AUC values even approached those for a double dose of nanoparticle without VDA (**Figure 2A, B**). The important difference between these groups is that the double dose also doubles systemic doxorubicin delivery e.g. in skin, heart, and lungs (**Figure 2C, D**; **Figure S2B**), whereas with VDA treatment, the enhancement is tumour selective.

**Figure 2.**
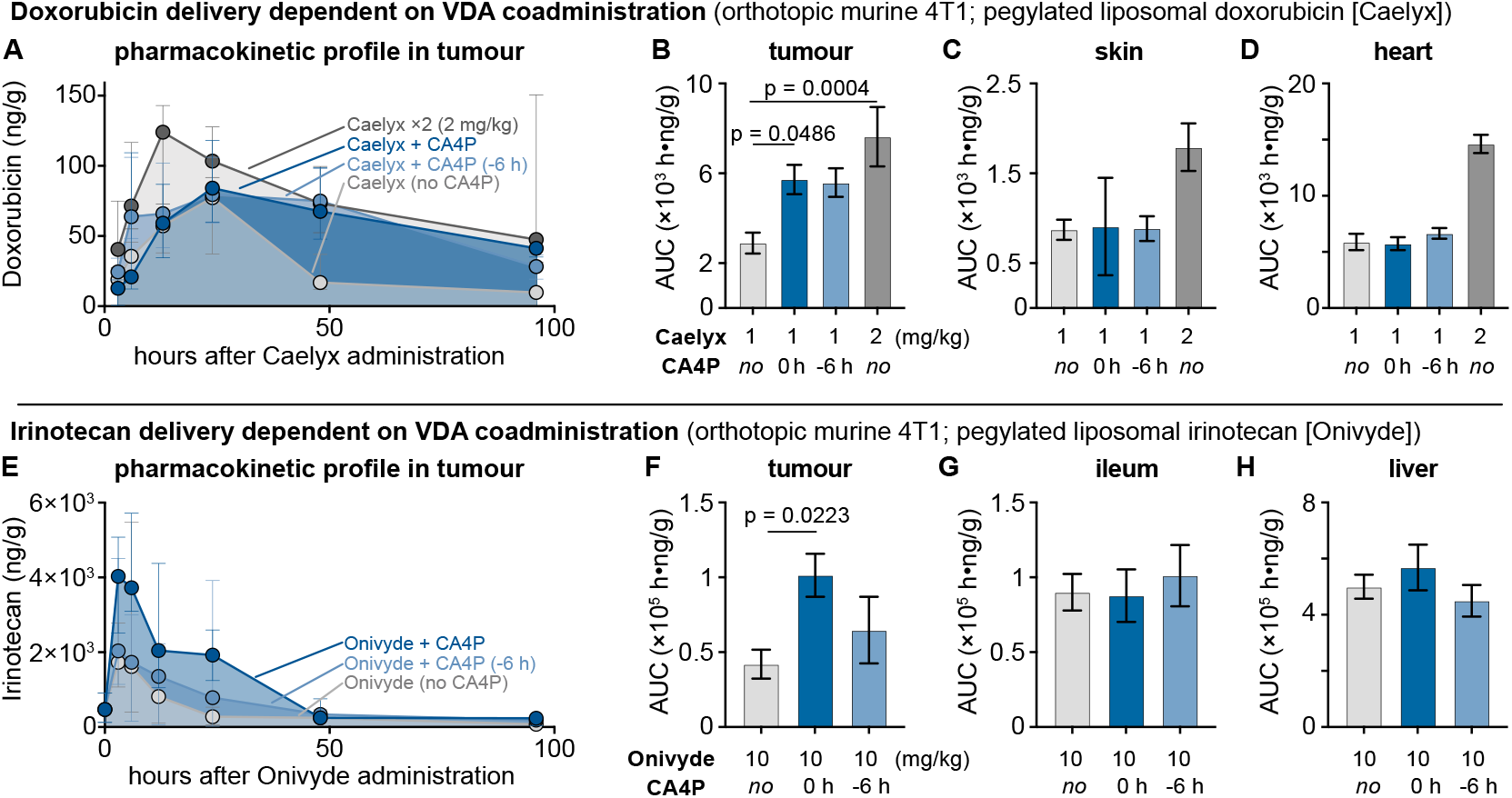
Co-administration of nano-formulated drugs with CA4P increases tumour exposure while sparing healthy tissue. LC-MS/MS quantification of doxorubicin (Caelyx®, 1 and 2 mg/kg) and irinotecan (Onivyde®, 10 mg/kg) in orthotopic 4T1 tumours and in healthy organs, from administering the nano-formulations either alone, or together with a CA4P dose (30 mg/kg), or with the CA4P dose having been given 6 h earlier (related data in **Figure S2**). (A, E) Tumour pharmacokinetics for doxorubicin and irinotecan. (B, F) Tumour AUC comparisons. (C, D) Skin and heart exposure for Caelyx groups. (G, H) Ileum and liver exposure for Onivyde groups. Median, 95% CI, n = 5 per time point, one-way ANOVA.

We also tested whether administering CA4P 6 hours before Caelyx might “prime” tumour vasculature for faster or increased delivery. Indeed, we measured a faster tumoral drug loading over the time window of 3 to 6 hours in the priming group, as compared to the co-administration group (**Figure 2A**). Still, the overall AUC values after 96 h were similar (**Figure 2B**), which is expected since Caelyx has a plasma circulation half-life ca. 18 h (**Figure S2A**), whereas VDA effects are transient and faster (half-life typically on a 6 hour timescale (Dark et al., 1997)) allowing both primed and co-administered groups to integrate similar vascular nanoparticle exposure during the long-lasting phase of highest systemic nanoparticle load (0-18 hours). For long-circulating diagnostic nanoparticles that should be imaged rapidly (e.g. PET reporters), such early-stage effects from VDA priming might be an important benefit; although for nanoparticles with shorter circulation times, or whose extravasation ability decreases rapidly by opsonisation, the time-dependency of priming effects might be complex to deal with.

#### Onivyde^®^ (irinotecan)

Coadministration of Onivyde (10 mg/kg) with CA4P more than doubled irinotecan AUC in tumour tissue compared to Onivyde alone (**Figure 2E, F**) and gave a five-fold AUC enhancement for its actual pharmacologically active metabolite species, SN-38 (**Figure S2E**). Free Onivyde is eliminated rapidly (half-life <3 hours, **Figure S2C**); faster tumoral loading and longer tumoral retention both contribute to this AUC enhancement. Crucially, CA4P did not increase irinotecan accumulation in organs commonly affected by off-target toxicity, including small intestine and liver (**Figure 2G, H**), nor in plasma or spleen (**Figure S2C, D**). This is important, as irinotecan-associated diarrhoea from intestinal mucosal damage often limits treatment intensity (Zhang, 2016).

Comparing Onivyde and Caelyx assays provided an opportunity to test the plausibility of the time-dependent effects we had expected to see, because the narrow time window for good Onivyde delivery, determined by high plasma availability of its nanoparticles (0-3 h, **Figure S2C**), does not match the apparent window for enhanced tumoral uptake in the VDA-primed group, determined by tumour vascular effects (3-6 h timepoints, see Caelyx study, **Figure 2A**). This gave the expectation that VDA priming 6 hours before Onivyde treatment might not effectively load tumours with irinotecan: and indeed, the primed group showed tumoral loading that was almost identical to that of the no-VDA group (**Figure 2E** light blue graph). Nonetheless, an overall tumoral AUC benefit driven by increased drug retention was still expected and observed for the priming group (**Figure 2F** light blue bar, ~1.5-fold AUC compared to no-VDA). While it might be beneficial to shift and align the VDA priming window to achieve tumoral distribution benefits depending on the pharmacokinetic properties for that specific nanoparticle, we consider this proof-of-concept test to be a robust result and decided to terminate biodistribution studies in favour of therapeutic assays.

Taken together, the Caelyx and Onivyde data are coherent with the reporter results (**Figure 1**), showing that VDA co-administration can selectively enhance the tumoral delivery and retention of nano-formulated chemotherapeutics, without increasing exposure in healthy organs. We now moved to assess whether it could be leveraged over multiple administration cycles to improve treatment outcomes, in the hope that tumour-enhanced nanoparticle delivery from VDA co-administration might allow e.g. more effective chemotherapy without increased systemic toxicity at current nanoparticle dosing, or equally effective chemotherapy with reduced systemic toxicity at lowered nanoparticle dosing.

### The tumour-selective delivery enhancement is not retained after the first treatment, and the VDA combination does not increase therapeutic efficacy in standard settings

Based on the tumour-selective increase in nano-formulated agents observed in the reporter and pharmacokinetic studies, we designed corresponding therapeutic studies in a syngeneic, immunocompetent murine model of triple-negative breast cancer (4T1).

#### Caelyx and Onivyde combination therapy with CA4P fails to improve tumour control

As expected, CA4P monotherapy at 30 mg/kg had little to no effect on tumour growth (**Figure 3A, 3B**, light red). Treatment with Caelyx at 2 mg/kg or Onivyde at 10 mg/kg slowed tumour progression (grey lines), consistent with their known efficacy in this model. However, co-treatment with CA4P did not further improve tumour control (blue lines). This absence of benefit was surprising given the initial increase in drug delivery we observed after a single co-administration. We hypothesised that this might reflect a loss of vascular susceptibility over time. Tumour blood vessels could remodel after repeated CA4P and drug exposure or become less permeable due to maturation or therapy-induced selection. To test this, we measured tumour drug levels after each treatment cycle.

**Figure 3.**
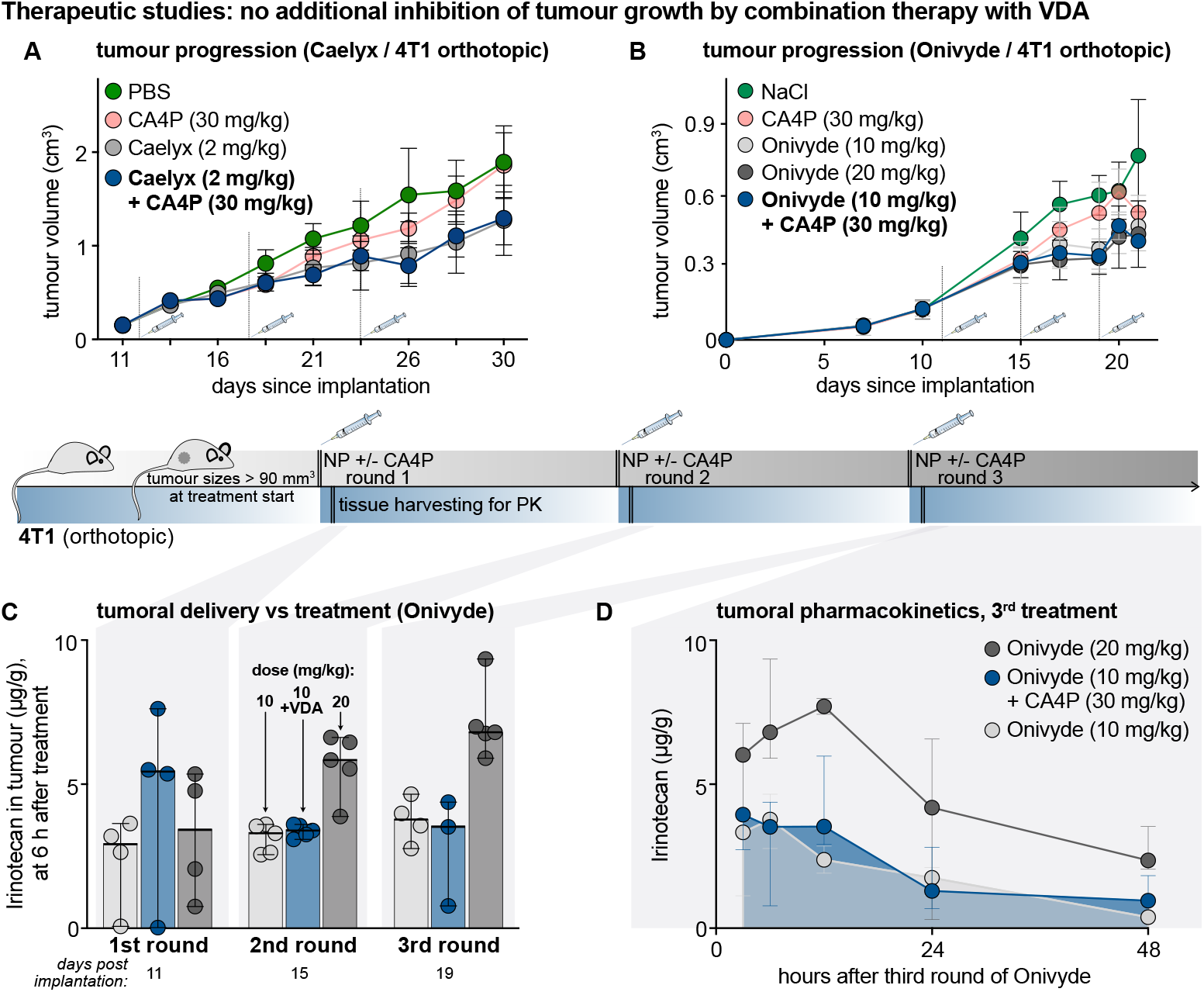
Limited therapeutic efficacy of CA4P in combination with nano-formulated chemotherapeutics. (A) Co-treatment with CA4P (30 mg/kg) and Caelyx (2 mg/kg) in the 4T1 breast cancer model (weekly dosing, n=5) does not improve tumour control compared to Caelyx monotherapy. (B) Co-treatment with CA4P (30 mg/kg) and Onivyde (10 mg/kg, every 4 days) shows no added benefit over Onivyde alone. Mean with SD, n ≥ 5 mice per group. (C) Irinotecan quantification in tumours at 6 h post-treatment shows increased accumulation only after the first combination dose. (D) Full PK analysis of irinotecan in tumours after three Onivyde treatments (10 or 20 mg/kg), with or without CA4P, shows no significant accumulation benefit from the combination. Median, 95% CI, n = 5 per time point.

#### CA4P-induced drug uptake enhancement is not maintained over multiple treatments

We analysed irinotecan levels in tumours at 6 hours after the first, second and third Onivyde treatment with and without CA4P (**Figure 3C**). While the first CA4P combination significantly increased irinotecan accumulation as previously shown (**Figure 2E, F**), this effect was lost by the second treatment. After the second and third cycle, tumour drug levels were indistinguishable between combination and monotherapy. A full PK profile after the third treatment confirmed this: the curve for combination treatment now matched monotherapy (**Figure 3D**), lacking the initial peak and extended retention seen after the first dose observed in **Figure 2E**. Conversely, a higher dose of Onivyde (20 mg/kg) that was included in this study as a control showed the expected doubling of tumour drug content compared to the standard dose after the second and third treatment (**Figure 3C**) and in the full third treatment PK curve (**Figure 3D**), although unexpectedly, no doubled accumulation during the first treatment round. Noteworthily, 20 mg/kg Onivyde did not show any therapeutic benefit over the 10 mg/kg Onivyde monotherapy or the combination therapy with CA4P (**Figure 3B**). We cannot exclude technical errors or biological variability contributing to this observation, although it may reflect a pharmacological difference to Caelyx, where a doubling of the dose produced a proportional increase in doxorubicin accumulation within the tumour (**Figure 2A**).

These results indicate that the window for CA4P to increase drug uptake via tumour vasculature is transient and most likely dependent on the maturity of the blood vessel system. Once structural vulnerability is lost, the combinations of Caelyx and Onivyde with CA4P behave no differently from the respective monotherapies. This likely explains the lack of improved tumour growth inhibition with either Caelyx or Onivyde when combined with CA4P. Due to animal welfare considerations, we did not repeat full PK profiling after multiple treatment cycles with Caelyx. However, given the consistency of previous data in terms of CA4P-mediated tumour selective drug accumulation in first round treatment, but a lack of therapeutic efficacy over multiple rounds, we would expect to find the same limited window of benefit. All animals included in efficacy studies bore tumours above 90 mm^3^ to ensure active neovascularisation (Carmeliet & Jain, 2000); **Figure S3**). Even so, vascular maturation during tumour growth, remodelling, and damage from prior treatments may reduce the efficacy of repeated CA4P vascular modulation.

## Conclusion and Discussion

Low-dose vascular disruption with CA4P can transiently and selectively increase the delivery of nano-formulated drugs to tumours without affecting healthy tissue. While the effect is not retained over repeated dosing, it provides proof-of-concept for vascular targeting as a delivery tool in research or diagnostic applications where a single, tumour-selective delivery pulse is sufficient.

Vascular targeting is a compelling route for improving drug delivery to solid tumours. Its potential for broad applicability across tumour types, without requiring tumour-specific ligands or immune phenotyping, stands in frugal contrast to personalised medicine strategies and antigen-based targeting. In this study, we tested the use of the vascular disrupting agent CA4P as a transient delivery enhancer, repurposing it from a therapeutic agent into a tool for improving intra-tumoral accumulation of nano-formulated chemotherapeutics. Complementary work on vascular disruption for agent delivery was previously described and our initial findings are in line with the work performed on the tumoral delivery of fluorescent dextran reporter molecules (Tozer, Ameer-Beg, et al., 2005), nano-formulated Paclitaxel (Gao et al., 2019), or radioisotope ^177^Lutetium complexes (Satterlee et al., 2017). This is the first time, however, that thorough long-term therapeutic and pharmacokinetic studies were performed with different nano-formulated, clinically approved therapeutic agents at low doses of CA4P and while monitoring drug accumulation in healthy tissue. We show that a single low dose of CA4P can increase drug delivery selectively to tumours, without increasing drug exposure in healthy organs.

Historically, vascular disrupting agents such as CA4P were developed as monotherapies to destroy immature tumour vessels and trigger necrosis (Siemann et al., 2009). Despite promising early-stage results, clinical trials showed limited efficacy and frequent recurrence, with viable tumour cells persisting in the well-perfused tumour margin. Combination strategies with other cytotoxic agents were largely unsuccessful, in part due to overlapping adverse effects. CA4P was associated with cardiotoxicity that restricted its clinical use. These limitations led us to shift focus from vascular targeting for therapy to delivery. We used subtherapeutic doses of CA4P to exploit the structural weakness of tumour neo vasculature and enhance nanoparticle delivery to tumours, while limiting systemic exposure.

Our findings confirm that this approach can transiently enhance drug accumulation in tumours. Both liposomal irinotecan and doxorubicin showed increased tumour uptake following the first dose of CA4P. Importantly, this effect was selective for tumour tissue, with no increase in drug accumulation observed in heart, skin, liver, spleen, kidney, or lung. The selectivity likely arises from structural differences between disorganised tumour vessels and stable vasculature in healthy organs. However, the delivery benefit did not translate into improved therapeutic efficacy.

We observed that the CA4P-mediated drug accumulation effect was not conserved beyond the first treatment cycle, most likely when immature vessels still dominate the tumour. Once these are lost, whether by VDA- or therapy-induced necrosis, natural maturation, or vascular remodelling, combinations of Caelyx or Onivyde with CA4P behave no differently from monotherapy. This loss of effect may reflect reactive vascular remodelling in response to therapeutic intervention or stabilisation of the vasculature over the course of tumour growth, meaning loss of structurally weak blood vessels as target for our delivery approach. Flow cytometry data from related experiments suggested that vessel maturity differs significantly between tumours of different growth kinetics, though we did not systematically explore these differences in the current study. Such plasticity in the tumour vasculature likely contributes to the inconsistency of the delivery effect.

The disconnect between increased drug delivery and therapeutic outcome also raises questions about the choice of drug and its mechanism of action. Liposomal formulations of topoisomerase inhibitors, which mainly target dividing cells during S-phase, so typically rely on repeated exposure for maximal efficacy, may therefore not be optimal candidates for a single-dose delivery enhancement strategy. Future work could explore more potent agents or substances with mechanisms of action better suited to short-term high-concentration delivery.

Our results also underscore the limitations of this approach in terms of reproducibility and model dependence. The delivery enhancement was highly variable across studies, even within the same tumour model. We conducted all pharmacokinetic and efficacy studies using orthotopic 4T1 tumours in immunocompetent mice, whereas imaging in Figure 1 was performed in subcutaneous 4T1 tumours in immunocompromised nude mice. Differences in tumour microenvironment, vascular architecture, and perfusion likely account for some of the variability observed. Only studies with valid controls were included in the final analysis. Nevertheless, the delivery effect likely proved sensitive to factors such as tumour size, location, vascularisation, and growth rate, many of which are difficult to control *in vivo* and even harder to predict in clinical settings.

Despite these limitations, the core finding remains robust: a single low dose of CA4P can produce a tumour-selective increase in nanoparticle delivery without increasing off-target exposure. Because the VDA effect seems independent of drug identity, future work might explore nano-formulations better suited to succeed with a one-time delivery boost. In exploratory assays, CA4P also increased tumour accumulation of nucleic acids (e.g. reporter DNA), suggesting possible applications where a single, antigen-independent delivery boost is sufficient, such as imaging or diagnostic tracer delivery.

In summary, we present a proof-of-concept for low-dose vascular disruption as a delivery-enhancing tool. While the effect is transient, model-specific, and difficult to control over multiple treatments, it offers a strategy for tumour-selective delivery without increasing systemic exposure. Given the narrow window of efficacy and high variability observed, we do not consider this approach ready for clinical translation in its current form. However, it may find applications in diagnostics or basic research, where a one-time delivery boost is sufficient and highly selective, antigen-independent targeting is desired.

## Materials and Methods

### Chemical Compounds

Caelyx® from Janssen and Onivyde® from Baxalata Innovations GmbH was purchased through the pharmacy of the LMU Munich University Hospital. Combretastatin-A4P disodium salt (CA4P) was purchased from SelleckChem (Catalog No. S7204; CAS No. 168555-66-6). SAIVI™ 715, PEGylated microspheres for small animal *in vivo* imaging were custom synthesised by ThermoFisher in the batches of different average sizes (40, 80, 110, 140, 160, 240, and 320 nm diameters, determined using Transmission Electron Microscopy). The nanoparticle core is surfactant-free white polystyrene latex (Cat. No. S37204, Molecular Probes™), doped with NIR fluorophore, and PEGylated using PEG MW 2000 end-capped with methoxy groups.

<u>Combretastatin A4 phosphate</u> (CA4P) is a water-soluble prodrug of combretastatin A4 (CA4), a potent colchicine-like tubulin binder that slows microtubule polymerisation and accelerates depolymerisation. Following administration, CA4P is rapidly dephosphorylated *in vivo* to its active form CA4, which binds to the colchicine site of β-tubulin.

<u>Caelyx^®^</u> is a liposomal formulation of doxorubicin hydrochloride, in which the drug is encapsulated in liposomes composed of fully hydrogenated soy phosphatidylcholine (HSPC), N-(carbonyl-methoxypolyethylene glycol 2000)-1,2-distearoyl-sn-glycero-3-phosphoethanolamine sodium salt (MPEG-DSPE), and cholesterol.

<u>Onivyde^®^</u> is a nano-liposomal formulation of irinotecan hydrochloride, in which the active drug is encapsulated in liposomes composed of distearoylphosphatidylcholine (DSPC), cholesterol, and a polyethylene glycol (PEG)-conjugated phospholipid, such as N-(carbonyl-methoxypolyethylene glycol 2000)-1,2-distearoyl-sn-glycero-3-phosphoethanolamine sodium salt (MPEG-DSPE).

### Animal husbandry

All animal experiments were conducted in accordance with EU animal welfare law (Directive 2010/63/EU on the protection of animals used for scientific purposes), the GV-SOLAS and FELASA guidelines and were approved by the appropriate local authorities. In Germany, experiments were authorised by Landesamt fuer Gesundheit und Soziales Berlin (LaGeSo), the Regional Board Freiburg (Regierungspräsidium) and the Regierung von Oberbayern, Munich (ROB Az. 55.2-2532.Vet_02-19-46). In France, the animal experiments and procedures complied with ethical rules for *in vivo* testing in mouse models and were approved by the relevant authority following French law to minimize animal suffering and the number of animals used. All animal experiments were approved by the Animal Ethics Committee of the French Ministry, under the agreement number APAFIS#8868-2015093015035547 v8 and were performed according to the Institutional Animal Care and Use Committee of Grenoble Alpes University. In Spain, experiments were approved by Comunidad Autonoma or Ministerio de Agricultura, Pesca y Alimentacion and reviewed by institutional Ethical Committee for Animal Experimentation (B-9900044 and A/ES/16/1-03; Notification date: 65-7951/2016).

BALB/c and NMRI nude mice were maintained in individually ventilated cages at constant temperature (22±2 °C, air-conditioned) and humidity (45 – 65%), under optimum hygienic conditions with 10 – 15 air changes per hour. A cycle of 12 h artificial fluorescent lightning and 12 h darkness was applied, and animal behaviour was monitored daily throughout the study. The mice received food and water *ad libitum*.

### Cell implantation, randomisation

4T1-luc2 cells were cultured in RPMI-1640 medium, supplemented with 10% FCS and penicillin 100 units/mL and streptomycin 100 µg/mL. 5×10^4^ 4T1-luc cells were injected in a mixture of 1/3 Matrigel and 2/3 RPMI (no supplementation) into the right flank of each mouse.

4T1 cells were cultured in RPMI-1640 high Glutamax with 10 % FCS, 100 units penicillin/mL and 100 µg streptomycin/mL. 10^5^ 4T1 cells in 100 µL PBS were implanted into the left mammary fat pad of each mouse.

Animals (BALB/c) were randomized after enough tumours had grown larger than 90 mm^3^, usually at a mean tumour volume of 200 – 250 mm^3^. Tumour volume was calculated as follows: V = 0.5 × length × width^2^

### Treatment

According to study design and treatment group, mice were injected with Onivyde^®^, Caelyx^®^ 4 mg/kg and/or CA4P or a vehicle of either PBS or 0.9 % NaCl. Onivyde^®^ and Caelyx were administered intravenously (i.v.), while CA4P was injected intraperitoneally (i.p.).

### Efficacy

After start of therapy, animal weight was determined three times weekly and tumour monitoring twice weekly by calipering. Deviation of the health status of the animals were documented, and animals were euthanized individually before study termination when ethical abortion criteria were reached (e.g. body weight loss ≥ 20%, signs of sickness; after s.c. or i.ma. implantation: tumour diameter ≥ 2 cm (s.c.), ≥ 1.8 cm (i.ma.), tumour ulceration). N.B. Several independent *in vivo* efficacy studies were conducted with Caelyx or Onivyde in combination with CA4P. Only studies in which the control treatments (Caelyx or Onivyde alone) showed consistent tumour growth inhibition, as expected from previously published and validated protocols, were included in the final analysis. Studies that failed to meet this internal benchmark were excluded from interpretation.

### Pharmacokinetics

After different numbers of treatment cycles, a set of 5 animals per experimental group was taken for necropsy at distinct timepoints (5 min, 3 h, 6 h, 13 h, 24 h, 48 h, 96 h). Blood was collected for the preparation of EDTA-plasma by retro-orbital vein puncture, centrifuged (4°C, 8000 rpm) and the supernatant was frozen at −80°C. Primary tumours and selected organs were collected, snap-frozen in liquid nitrogen and stored at −80°C until analysis.

### *In vivo* imaging

Female NMRI nude mice (6 weeks old, JANVIER) received a subcutaneous (s.c.) xenograft of 4T1-luc2 cells (ATCC^®^ CRL-2539™, 1✕10^6^ in 50 µL) in the right flank. After implantation tumour volume was measured three times per week using callipers. The tumour volume was calculated as follows: length × (width)2 ✕0.4. At post-implantation day 11 tumours would typically reach volume ~ 250 mm^3^. Mice were monitored and weighed three time per week. After 72 h post injection of drug and nanoparticles the mice were euthanized and tumours and selected organs isolated. Tissue was snap frozen and kept at −80°C until analysis. CA4P and nanoparticles were intravenously co-injected into the tail vein. NPs were checked for linearity of fluorescence with increased dilution. Since NPs intravenous injection in mice induces an immediate dilution in blood (~1/10) NPs solutions were used at ½ dilution for experimentation in mice.

Whole body *in vivo* fluorescence imaging (lateral side, right and left) was carried out in a Fluobeam700 (Fluopics), by using the respective wavelengths for excitation (680 nm) and emission (>700 nm). Each animal was imaged at timepoints t = 0, 6 h, 24 h and 48 h. Mice were sacrificed 48 h after treatment and selected organs were isolated for *ex vivo* fluorescence imaging.

In addition, ex vivo fluorescence imaging was performed on isolated organs at 72 h post-injection in heart, lung, brain, liver (right median lobe), spleen, left kidney, bladder, adrenal glands, jejunum, muscle (quadriceps), skin fragment, sexual organs (uterus and ovaries), pancreas, abdominal fat.

### LC-MS/MS

All LC-ESI-MS/MS measurements were performed on an API5000 triple quadrupole mass spectrometer with TurboV-ESI source (Sciex, Darmstadt, Germany). An Agilent 1200 Series HPLC system (vacuum degasser G1379B, binary pump G1312B, oven 1316B, Agilent, Waldbronn, Germany) and an HTS-PAL auto sampler (CTC-Analytics, Zwingen, Switzerland) was coupled to the mass spectrometer and Analyst v. 1.6.1 software (Sciex, Darmstadt, Germany) was used for hardware control, as well as for Data analysis.

#### Tissue Extraction

To extract the respective analytes of interest after Caelyx or Onivyde treatment, tumours, plasma and organs were homogenized in 5x volume w/v ethanol or acetonitrile, in a MP FastPrep™24-5G tissue-cell homogenizer (8 m/s, 30-60 s). The supernatants were collected after centrifugation at 13300 rpm for 5 min and stored at −80 °C until analysis.

Calibration curves were obtained by diluting ethanolic stock solutions (250 µM, Doxorubicin, Doxorubicinol) or methanolic stock solutions (10 µM, Irinotecan, SN-38, Camptothecin) with the corresponding extracts from blank tumours, tissue or plasma samples. Calibration curves for quantifications were established at concentrations of 1000 nM, 750 nM, 500 nM, 250 nM, 100 nM, 50 nM, 25 nM.

Internal standards were used at a constant final concentration of 50 nM per sample, whereby Daunorubicin was used for measurements of Doxorubicin and Doxorubicinol, and Camptothecin for the quantification of Irinotecan and SN-38. For analysis, 150 µL of each sample extract was mixed with 50 µL 200 nM internal standard solution and 10 µL of each mix were injected into the LC-MS/MS.

#### HPLC Methods

Both compound sets, derived from Caelyx or Onivyde, were analysed on a YMC-Triart PFP column (50 × 2.0 mm, 3 µm), with two inline filters (0.5 and 0.2 µm) installed before the column. The mobile phase composition varied slightly for the relevant two sets, however both relying on A: ammonium acetate (5 mM, pH = 3.5) and B: acetonitrile. For the analysis of Doxorubicin, Doxorubicinol and the internal standard Daunorubicin, a composition of 60:40 v/v (A:B) with a flow rate of 400 µL/min was applied. Irinotecan, SN-38 and the internal standard Camptothecin were analysed with a mobile phase of 63:37 v/v (A:B) and a flow rate of 350 µL/min. If not indicated differently 10 µL were injected per sample.

#### MS/MS parameters

For the determination of all compound-dependent MS parameters, a 20 nM solution of each analyte (Irinotecan, SN-38, Camptothecin, Doxorubicin, Doxorubicinol, Daunorubicin) in mixture of methanol and 0.1% formic acid, was infused directly into the ESI source, via an external syringe pump with a flow rate of 10 µL/min. Samples were acquired in Multiple Reaction Monitoring (MRM), positive mode with an ionspray voltage (IS) of 4500 V at 650 °C (curtain gas 25 psi, gas 1 55 psi, gas 2 45 psi).

Parameters determined by compound optimisation mode of the Analyst software, as well as compounds’ mass transitions, are available in the supporting information (table 1 and table 2).

### Fluorimetry

Extraction of the NIR reporter fluorophore from isolated tissue from in vivo imaging studies was performed as follows. Frozen tumour or tissue samples were homogenised using a FastPrep-24™ 5G (MP) and subdivided into two aliquots per tumour. Tissue homogenates were centrifuged briefly in a tabletop centrifuge and resuspended in 100 µL of 0.9% NaCl followed by the addition of 300 µL of *n*-butyl acetate. Samples were mixed on an Eppendorf tube rack overnight and centrifuged at 13000 × g for 10 min to separate phases. The upper organic phase, containing the extracted fluorophores, was carefully collected (60 µL) into acetone-rinsed microcuvettes and fluorescence was measured at 735 nm. Cuvettes were washed with acetone three times between measurements to prevent cross-contamination.

### Statistics

All statistical analyses were performed using GraphPad Prism version 8.4.2. Data are presented as mean or median with 95% confidence intervals, as indicated in figure legends. For pharmacokinetic analyses area under the curve (AUC) was calculated using the trapezoid rule. Statistical significance was assessed using ordinary one-way ANOVA followed by Dunnett’s multiple comparisons test, comparing each treatment group to a control. Statistical analysis of tumour growth in efficacy studies was performed using two-way repeated-measures ANOVA with Sidak’s multiple comparisons test in GraphPad Prism v10. A p-value less than or equal to 0.05 was considered statistically significant (alpha = 0.05).

## Acknowledgements

We thank Monique Preusse and the LMU Core facility for flow cytometry for excellent technical support. We thank Bettina Stahnke and team at Reaction Biology (Freiburg, Germany); Ramon Messeguer and team at Leitat (Barcelona, Spain); Diana Behrens and team at EPO (Berlin, Germany); and Anna Kondratiuk, Yuliia Holota, Sergey Zozulya, and the other team members at Bienta (Kyiv, Ukraine), for *in vivo* proof of concept and confirmatory studies, and *in vivo* PK studies. O.T.-S. thanks the German Research Foundation (DFG) for position support through an Emmy Noether grant (number 400324123). J.T.-S. thanks the Joachim Herz Foundation and the LMU Center for Nanoscience for support. The graphical abstract was created in https://BioRender.com

## Funding

This research was supported by the German Federal Ministry of Education and Research BMBF through Discovery (grant number 031B0159), Feasibility (grant number 031B0396), and GO-Bio (grant numbers 031B0632 and 161B0632) grants to O.T.-S., P.M., and J.T.-S.. The Optimal small animal imaging platform is supported by France Life Imaging (French program “Investissement d’Avenir” grant; “Infrastructure d’avenir en Biologie Santé”, ANR-11-INBS-0006) and the IBISA French consortium “Infrastructures en Biologie Santé et Agronomie”.

## Author Contributions

*In vivo* studies (imaging and PK) were performed by A.K., C.H., S.S., J.V., and J.T.-S., with supervision and coordination by V.J., J.-L.C., and J.T.-S. Drug and fluorophore extractions followed by instrumental analysis were performed by A.K., C.H., S.S., D.B., G.H., and J.T.-S. Cell and tissue biology experiments were conducted by A.K., C.H., S.S., J.H., D.B., M.R. and J.T.-S. Biochemical and analytical method development were conducted by A.K., G.H., and O.T.-S. Data analysis and interpretation were carried out by A.K., O.T.-S., P.M., and J.T.-S. Project administration and funding were provided by A.K., K.K., O.T.-S., P.M., and J.T.-S.. Coordination and supervision of all experiments, data assembly, and visualization, were conducted by J.T.-S. The manuscript was written by J.T.-S., with A.K. and O.T.-S., and input from all authors.

## Notes

### Competing Interest Statement

The authors have declared no competing interest.

