## Supporting Information for "The delivery of nano-formulated drugs to solid tumours is selectively increased by co-application of the vascular disrupting agent CA4P"

[1] Department of Pharmacy, LMU Munich, Munich, Germany, [2] Grenoble Alpes University, INSERM U1209, CNRS UMR5309, Institute for Advanced Biosciences, Cancer Targets and Experimental Therapeutics team, 38000 Grenoble, France, [3] Faculty of Chemistry and Food Chemistry, Dresden University of Technology, Dresden, Germany, [4] Institute for Clinical Chemistry and Laboratory Medicine, University Hospital and Faculty of Medicine, Technical University Dresden, Dresden, Germany.

***

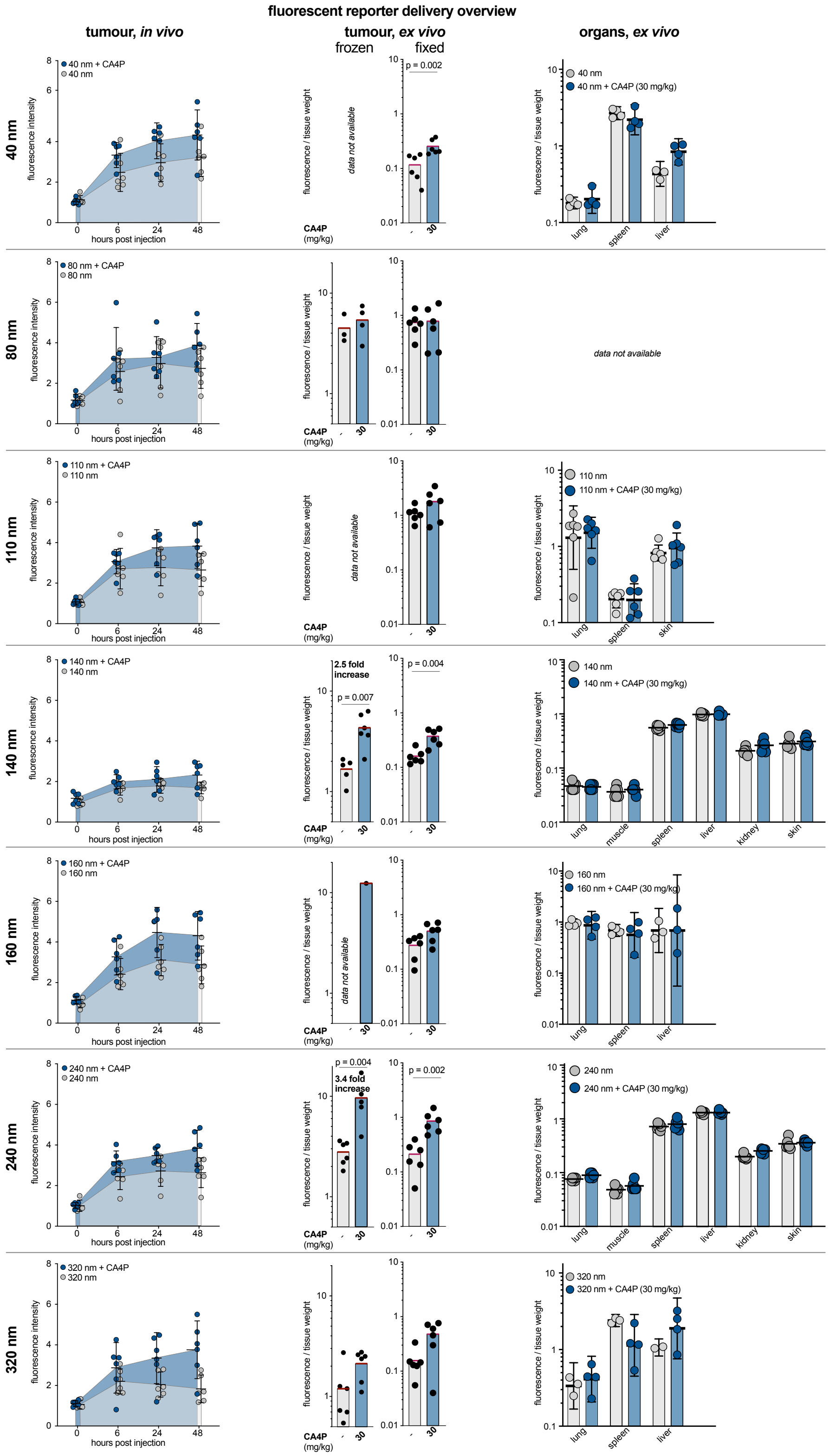


**Figure S1**: Tumour/tissue distribution of the NIR fluorophore reporter contained in SAIVI nanoparticles ("small animal in vivo imaging" nanoparticles, ThermoFisher), administered to mice with subcutaneous, syngeneic, triple-negative breast cancer (4T1) tumours, with (blue) or without (grey) CA4P. Reporter biodistribution was assessed qualitatively through longitudinal *in vivo* imaging, then quantitatively at the final imaging time point by sacrifice, *ex vivo* extraction of the fluorophore from snap-frozen and/or fixed tissues, and fluorimetric quantification normalised to tumour/organ weight. The nanoparticles tested covered a size range from 40 to 320 nm (40, 80, 110, 140, 160, 240, 320 nm), with the same interior composition and surface chemistry. Data were acquired over multiple experiment rounds; assays were not performed ("data not available") if initial rounds showed insufficient enhancement of accumulation.

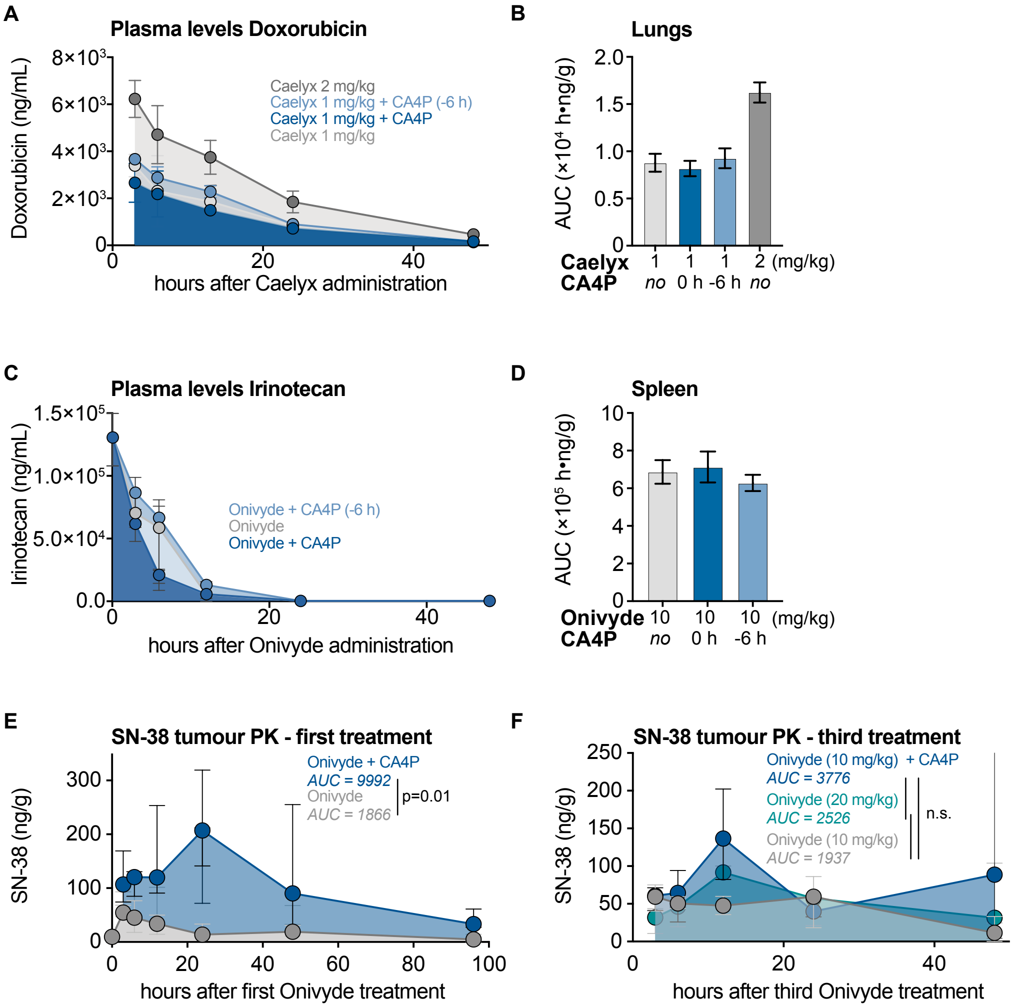


**Figure S2: Plasma pharmacokinetics and organ AUCs for Caelyx and Onivyde with or without CA4P.**
Plasma concentrations of doxorubicin (Caelyx, A) and irinotecan (Onivyde, C) were measured over 48 hours post administration, with and without co-treatment with CA4P. Corresponding area under the curve (AUC) values were calculated for doxorubicin in lung tissue (B) and for irinotecan in spleen (D) under the same treatment conditions. Note that the results from 1 vs 2 mg/kg doxorubicin groups, here and in **Figure 2**, display a proportional rise in both tumour *and* other organ drug levels: confirming both the analytical dynamic range and absence of saturation in the model (further details in the LCMS/MS *Methods* section). (E, F) Quantification of SN-38 (active metabolite of irinotecan) in tumours by LCMS/MS showing significant increase in AUC with CA4P after first treatment, but no CA4P-mediated significant increase after third treatment.

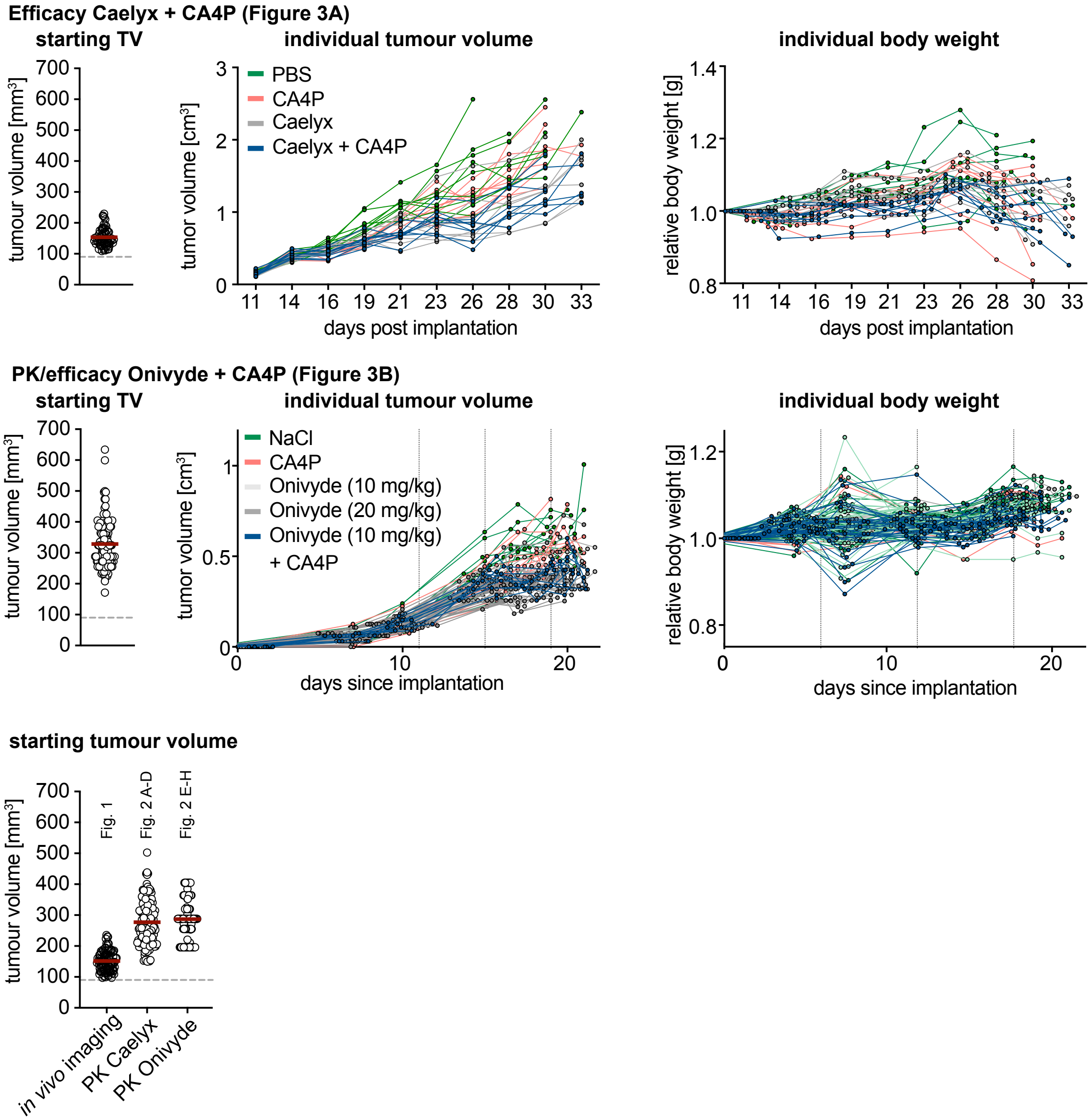


**Figure S3: Tumour volumes at treatment starts, individual growth curves and body weight tracking.** Starting tumour volume distributions for all included studies are shown. Individual tumour growth curves and corresponding body weights were tracked throughout the therapeutic study duration for each treatment group and mouse.

### MS/MS parameters for Irinotecan, SN-38, Camptothecin, Doxorubicin, Doxorubicinol, Daunorubicin

For the determination of all compound-dependent MS parameters, a 20 nM solution of each analyte in mixture of methanol and 0.1% formic acid, was infused directly into the ESI source, via an external syringe pump with a flow rate of 10 µL/min. Samples were acquired in Multiple Reaction Monitoring (MRM), positive mode with an ionspray voltage (IS) of 4500 V at 650 °C (curtain gas 25 psi, gas 1 55 psi, gas 2 45 psi).

By using the compound optimisation mode of the Analyst software, the following parameters were determined for each compound:

Table 1: Compound dependent MS/MS parameters for Doxorubicin and related, physicochemically similar species used as calibrants and standards (Doxorubicinol, Daunorubicin, Idarubicin). Q1= $[M+H]^+$ parent ion, Q3=fragment ions, EP= entrance potential, DP= declustering potential, CE= collision energy, CXP= cell exit potential

| compound | Q1 | Q3 | Time [ms] | EP [V] | DP [V] | CE [V] | CXP [V] |
|---|---|---|---|---|---|---|---|
| **Doxorubicin** | 544.4 | 483.4 | 150 | 10 | 86 | 17 | 50 |
| | | 397.2 | 150 | 10 | 86 | 33 | 14 |
| | | 527.5 | 150 | 10 | 86 | 21 | 18 |
| **Doxorubicinol** | 546.5 | 399.1 | 150 | 10 | 101 | 21 | 20 |
| | | 363.2 | 150 | 10 | 101 | 15 | 14 |
| | | 529.4 | 150 | 10 | 101 | 19 | 20 |
| **Daunorubicin** | 528.4 | 321.3 | 150 | 10 | 86 | 41 | 22 |
| | | 381.2 | 150 | 10 | 86 | 13 | 14 |
| | | 363.0 | 150 | 10 | 86 | 21 | 32 |
| **Idarubicin** | 428.3 | 291.1 | 150 | 10 | 76 | 45 | 44 |
| | | 351.1 | 150 | 10 | 76 | 13 | 54 |
| | | 333.1 | 150 | 10 | 76 | 23 | 6 |

Table 2: Compound dependent MS/MS parameters for inactive prodrug Irinotecan, its active metabolite SN-38, and physicochemically similar species Camptothecin (used as calibrant standard). Q1= $[M+H]^+$ parent ion, Q3=fragment ions, EP= entrance potential, DP= declustering potential, CE= collision energy, CXP= cell exit potential

| compound | Q1 | Q3 | time | DP | EP | CE | CXP |
|---|---|---|---|---|---|---|---|
| **Irinotecan** | 587.3 | 195.3 | 150 | 246 | 10 | 39 | 12 |
| | | 167.2 | 150 | 66 | 10 | 57 | 18 |
| **SN-38** | 392.2 | 315.0 | 150 | 66 | 10 | 21 | 22 |
| | | 349.0 | 150 | 66 | 10 | 39 | 12 |
| **Camptothecin** | 349.1 | 305.3 | 150 | 66 | 10 | 33 | 34 |
| | | 271.0 | 150 | 66 | 10 | 13 | 20 |
